# Designed Nanoparticles Elicit Cross-Reactive Antibody Responses To Conserved Influenza Virus Hemagglutinin Stem Epitopes

**DOI:** 10.1101/707984

**Authors:** Dustin M. McCraw, Mallory L. Myers, Neetu M. Gulati, John R. Gallagher, Alexander J. Kim, Udana Torian, Audray K. Harris

## Abstract

Despite the availability of seasonal vaccines and antiviral medications, influenza virus continues to be a major health concern and pandemic threat due to the continually changing antigenic regions of the major surface glycoprotein, hemagglutinin (HA). One emerging strategy for the development of more efficacious seasonal and universal influenza vaccines is structure-guided design of nanoparticles that display conserved regions of HA, such as the stem. Using the H1 HA subtype to establish proof of concept, we found that an alpha-helical fragment (helix-A) from the conserved stem region can be displayed on nanoparticles. The stem region of HA on these nanoparticles is immunogenic and the nanoparticles are biochemically robust in that heat exposure did not destroy the particles and immunogenicity was retained. Furthermore, H1-nanoparticles protected mice from lethal challenge with H1N1 influenza virus. Importantly, antibodies elicited by these nanoparticles demonstrated homosubtypic and heterosubtypic cross-reactivity. The helix-A stem nanoparticle design represents a novel approach to display several hundred copies of non-trimeric conserved HA stem epitopes on vaccine nanoparticles. This design concept provides a new approach to universal influenza vaccine development strategies and opens up opportunities for the development of nanoparticles with broad coverage over many antigenically diverse influenza HA subtypes.

**Significance:** Influenza virus is a public health issue that affects millions of people globally each year. Commercial influenza vaccines are based on the hemagglutinin (HA) surface glycoprotein, which can change antigenically every year, demanding the manufacture of newly matched vaccines annually. HA stem epitopes have a higher degree of conservation than HA molecules contained in conventional vaccine formulations and we demonstrate that we are able to design nanoparticles that display hundreds of HA stem fragments on nanoparticles. These designed nanoparticles are heat-stable, elicit antibodies to the HA stem, confer protection in mouse challenge models, and show cross-reactivity between HA subtypes. This technology provides promising opportunities to improve seasonal vaccines, develop pandemic preparedness vaccines, and facilitate the development of a universal influenza vaccine.

## Introduction

Influenza virus infects millions of people worldwide every year and there is the ever-present threat of a pandemic like the 1918 influenza virus that killed millions (1). Vaccines are formulated with influenza glycoproteins (2-5) and elicit antibodies to hemagglutinin (HA), the major surface glycoprotein (6-8). However, current commercial influenza vaccines must be manufactured each year with the assertation that elicited antibodies will provide immune responses to influenza viruses that are predicted to be circulating in an upcoming influenza season. In addition, these vaccines have limited immune responses to different influenza viral strains and subtypes that could enter the human population (9, 10). HA molecules for influenza A viruses exist as antigenically distinct subtypes ranging from H1 to H18 (11) within two phylogenetically-distinct groups (group 1 and group 2). Within these two groups, virus subtypes like avian H7N9, that have not circulated widely in humans, represent potential pandemic threats (12). One approach to preparing for a pandemic outbreak is to establish stockpiles of vaccines for high-risk pandemic threats. However, the constant antigenic drift of HA and the limited shelf-lives of conventional vaccines make them not conducive to stockpiling and millions of doses of influenza vaccines must still be produced annually to replace the expired vaccines (13-15).

All influenza HA subtypes have a similar protein structure (16, 17). HA exists as a protein trimer, produced as a single protein termed HA0, and is cleaved into disulfide-linked HA1 and HA2 proteins (18, 19). HA1 forms an apical globular region referred to as the head region. HA2 forms the stem region, along with some extended regions of HA1. While antibodies can bind to the head and stem of influenza, HA1 is immunodominant and more immunogenically variable than the more conserved HA2 stem region (20, 21). The HA head region appears to evolve faster than the stem region (22). This necessitates that HA vaccines be reformulated every year to match the continually changing head epitopes of circulating viruses. This explains why the estimated effectiveness of commercial vaccines can vary and can be less than 50% in some years (23).

One global goal in public health is the development of a universal influenza vaccine (24, 25) that could provide broad immunity to different strains and subtypes of influenza (10, 26). The ability to stockpile such a vaccine would aid in pandemic preparedness. One emerging strategy for the development of a universal influenza vaccine has been to focus on producing antigens, such as engineered proteins and nanoparticles, that elicit antibodies to conserved epitopes of HA and lead to broader protection (27-35). It has been speculated that this may require an iterative approach to antigen design and evaluation in order to develop a more efficacious vaccine (24). Due to the immunodominance and variability of the HA head region, one emerging concept is to design immunogens that display conserved HA stem regions. Several antibodies that bind to HA stem regions, termed stem antibodies, have been identified and shown to provide protection against influenza challenge (28, 36-45). The epitopes of some of these antibodies, including human monoclonal antibodies FI6V3 and CR6261, have also been structurally mapped (17, 28, 30, 36, 37, 46-48). This has facilitated interest in structure-guided efforts to engineer trimeric HA2 stem immunogens that elicit stem antibodies (29, 35, 49-52). However, HA2 stem trimers require multiple targeted mutations to stabilize the mapped stem epitopes found on the prefusion structure of trimeric HA2 and these mutations vary based on the influenza subtype. Additionally, the majority of epitopes for the stem region of HA are present on HA2, but small segments of HA1 also contribute to some epitope footprints. This may complicate stabilization strategies and hinder implementation for developing HA2 trimer immunogens to all strains, subtypes and types of influenza virus. The trimeric stem regions also contain glycosylation sites that may interfere with antibody access. To avoid these complications, we postulated that a smaller stem epitope footprint that is non-trimeric may be advantageous in increasing multivalent display, stability, immunofocusing, and a broader applicability of HA stem-based immunogen design to different influenza A virus subtypes.

In this work, we designed a novel HA-stem nanoparticle platform using structure-guided techniques. Bioinformatics was used to identify a conserved peptide fragment of influenza, known as helix-A, located within the stem epitope footprint of several broadly neutralizing stem antibodies. Chimeric constructs were designed to place this HA helix A into the immunodominant loop of the hepatitis B virus (HBV) capsid, thereby using the capsid as a nanoparticle scaffold. Using helix-A from pandemic H1N1 influenza virus as a proof of concept, we designed and purified H1-nanoparticles and used structural methods to characterize the particles. We found that these nanoparticles were immunogenic in mice and elicited antibodies capable of binding full length H1 HA protein. Immunization with the H1-nanoparticles protected mice from lethal challenge with H1N1 virus. Also, HA stem nanoparticles could be produced for HA representing both group 1 and group 2 influenza A viruses. Furthermore, we found that the HA nanoparticle antibody sera exhibited both homosubtypic and heterosubtypic binding activity to H1 and H7 HA proteins. These results establish proof of concept for the development of a nanoparticle-based vaccine that displays helix-A of the influenza HA stem region. This new vaccine nanoparticle platform will facilitate the design of multiple antigens that could open new opportunities for the production of more efficacious seasonal and pandemic vaccines, and facilitate universal influenza vaccine development.

## Results

### Helix-A epitope library sequence selection

The stem region of HA contains epitopes for a number of antibodies, some of which have been shown to bind to HAs from multiple subtypes (30, 36, 47, 53) (e.g. Fig. S1A-C). There is an accessible alpha helix within the HA2 stem region called helix-A, also called the A-helix (Fig. 1A-1C). The epitopes of several well characterized stem-binding antibodies include the helix-A region of HA2, making it an interesting epitope target for a vaccine candidate (Fig. S1). To facilitate the design of nanoparticles that integrate helix-A, we used bioinformatics techniques to assess the sequence conservation between HA subtypes. Primary amino acid sequences (N=50,428) for HA molecules from influenza type A virus were downloaded from the influenza sequence database (fludb.org) for H1-H16 HA subtypes. The helix-A region, which contains 22 residues, was then isolated from the HA sequences. Helix-A consensus sequences representing HA subtypes were found to be conserved and ranged from 50% to 100% sequence identities between HA subtypes (Fig. S2). For example, helix-A sequences from H1 and H2 proteins have 86% sequence identity, while H4 and H14 subtypes have 100% sequence identity within the helix-A stem epitope (Fig. S2).

**Fig. 1.**
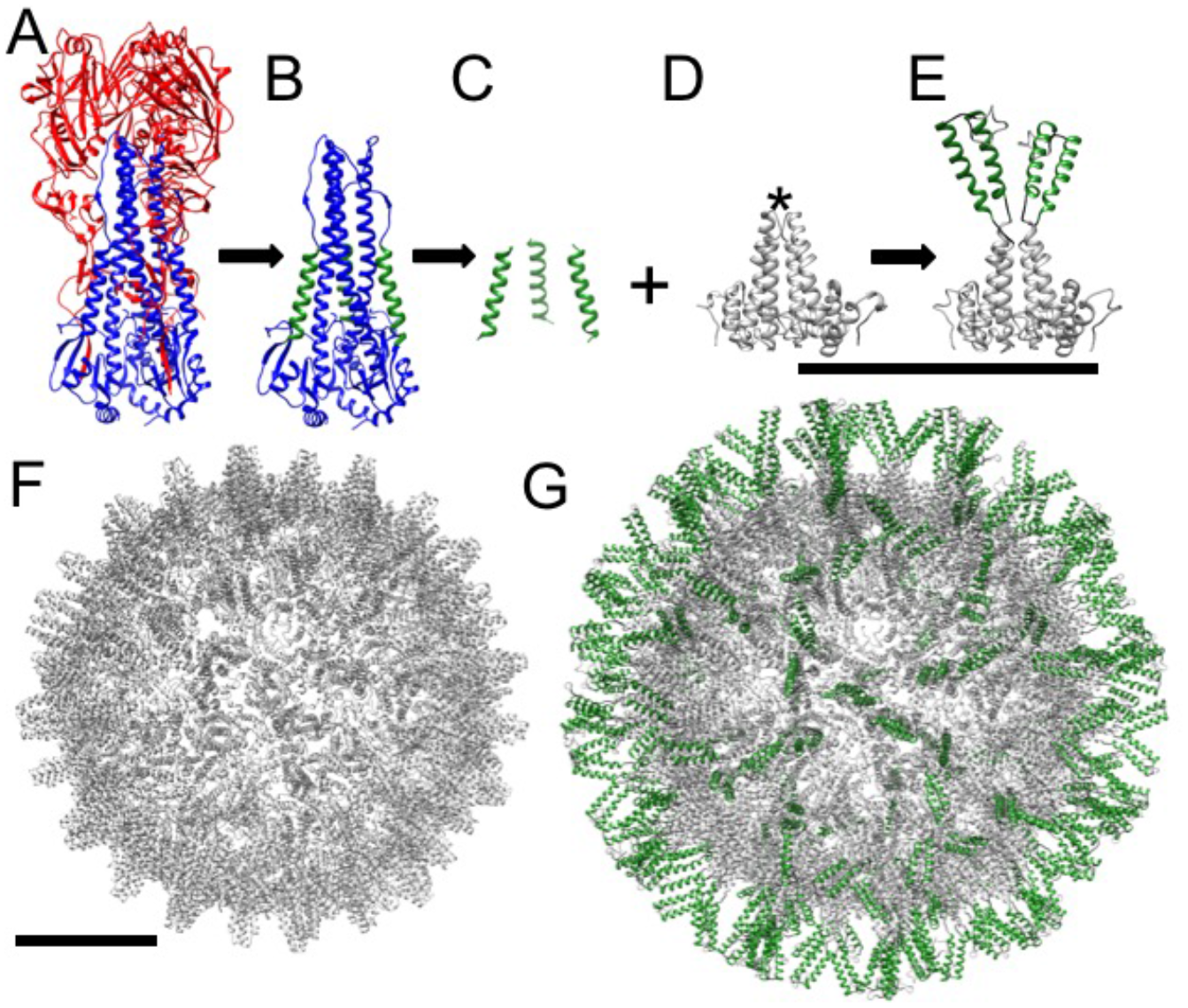
Nanoparticle design integrating the conserved helix A of hemagglutinin (HA) onto a nanoparticle scaffold. (A) Structure of influenza H1 HA ectodomain (PDBID 3LZG). HA1 is shown in red and HA2 in blue. (B) HA2 with the HA1 region computationally removed. The conserved helix A is shown in green within the HA2 stem region. (C) Computationally extracted helix-A segments are shown. (D) The scaffold, hepatitis B virus (HBV) capsid dimer, showing the alpha helical fold of the protein. The immunodominant loop at the tip of the dimeric spike is denoted by an asterisk, which is also referred to as the c1 epitope. (E) Homology model of the designed H1-nanoparticle dimeric unit. The protein design consists of a capsid monomer with two copies of helix A (green) inserted into the tip of the loop of the capsid protein (gray). Scale bar is 10 nm. (F). Structure of the scaffold (HBV capsid (PDBID 1QGT)), which has T=4 icosahedral symmetry and 240 dimer subunits per capsid. (G) Homology model for the H1-nanoparticle with icosahedral symmetry. The surface has the helix-A stem epitopes of HA (green) on the surface with the capsid scaffold (gray) forming the base core of the nanoparticle. Scale bars 5 nm.

Due to the conservation between helix-A epitopes from different HA subtypes, we selected a library of 27 sequences to represent HA sequence diversity. The library contains one or more helix-A sequences from each HA (H1-H16) subtype (Fig. S3A). Bioinformatics analyses indicated that the library of helix-A sequences matched greater than 91% of reported HA sequences to 95% identity or better (Fig. S3B). Taken together, these data suggest that the sequence library comprehensively represents the conserved helix-A stem region from most HA subtypes.

### Nanoparticle design

To probe whether nanoparticles that display the helix-A sequence could be expressed, we designed nanoparticles that integrated sequences from our helix-A library into the exposed immunodominant loops of the HBV capsid scaffold (Fig. 1D asterisk, 1E). We chose this scaffold because it is known to be temperature-stable (54) and amenable to sequence insertions (55). The scaffold itself is composed of 240 monomeric units which form dimers that assemble into an icosahedral capsid of approximately 30 nm (Fig. 1F). Our design parameters integrated two copies of helix-A into each monomeric unit, (Fig. 1E, S4) which is equivalent to 480 copies of helix-A sequences per nanoparticle (Fig. 1G). We hypothesized that the helix-A insertions would be displayed on the surface of the nanoparticle based on the scaffolding structure, and furthermore it would project from the immunodominant region of the capsid. Therefore, it would be likely to elicit antibodies to the helix-A insertion. To facilitate nanoparticle folding, we integrated flexible linkers flanking the helix-A insertions (Fig. S4).

Homology modeling was used to predict the structure of the helix-A insertion on the nanoparticle scaffold, using an integrated helix-A sequence from the H1N1 2009 pandemic influenza virus (A/California/07/2009) as a proof of concept. One representative homology model shows the helix-A insertion approaching a vertical orientation (Fig. 1E). Other representative homology models show the helix-A insertion approaching horizontal orientations (Fig. S5A). When icosahedral symmetry was applied to these models, all but one homology model displayed the helix-A epitopes protruding outwards from the surface of a successfully formed nanoparticle (Fig. 1G, S5B). In these nanoparticle constructs, the scaffold remained relatively unchanged, which suggests the insertion of the helix-A epitope does not hinder nanoparticle formation. Furthermore, while the scaffold is seemingly stable, the inserted epitope is likely flexible and may adopt multiple orientations.

### H1 nanoparticle production and purification

As a proof of concept for our helix-A nanoparticle design, we produced and purified nanoparticles with helix-A from the H1N1 2009 pandemic influenza virus (A/California/07/2009). This virus caused a pandemic outbreak of influenza in 2009 and since then, the circulating H1N1 influenza virus has been pandemic-like (pdm09-like). Both scaffold and designed H1-nanoparticle proteins could be expressed and purified as nanoparticles (Fig. S6). Purification of H1-nanoparticles by gradient centrifugation resulted in a banding pattern corresponding to a refractive index of 1.37, suggesting particles in excess of megadalton size (Fig. S7A-7D). The purity of the H1-nanoparticle (H1-nano) was observed to be greater than 95%, as determined by SDS-PAGE (Fig. 2A). Negative-stain electron microscopy confirmed the presence of nanoparticles of approximately 30 nm in diameter, which correlates to the observed gradient banding pattern (Fig. 2B, S7).

**Fig. 2.**
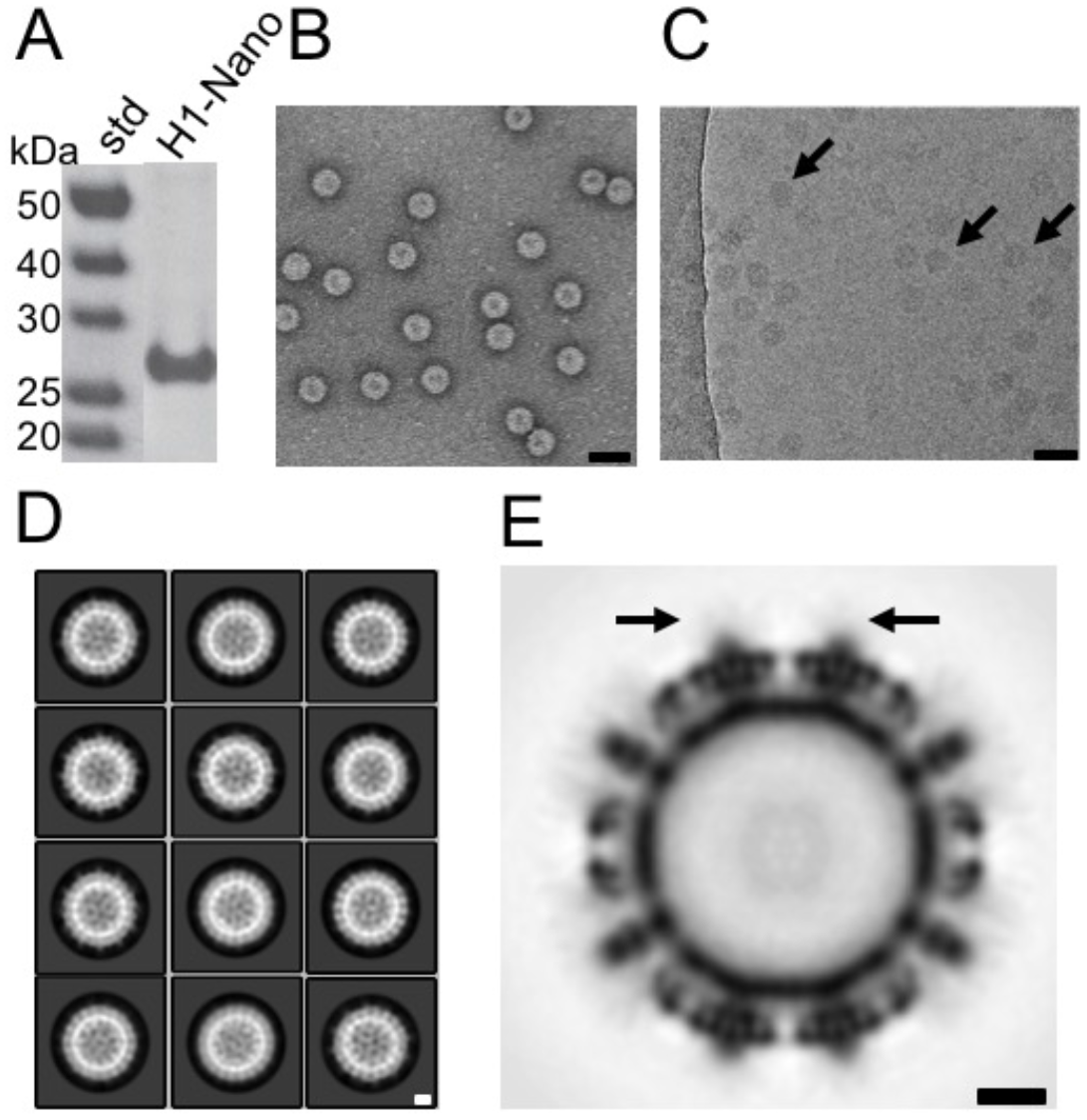
Structural analysis of purified H1-nanoparticle by electron microscopy and 3D reconstruction. (A) Analysis of purified H1-nanoparticle protein (H1-Nano) by SDS-PAGE with standards (std). (B) Image from negative-staining electron microscopy of the purified H1-nanoparticles. Scale bar 50 nm. (C) Cryo-electron microscopy image of a field of H1-nanoparticles. Arrows denote particles. Protein is black. Scale bar 50 nm. (D) Reference-free 2D class averages of the H1-nanoparticle. Protein is white. Scale bar 5 nm. (E) Central-slice through the 3D reconstruction of the H1-nanoparticle. Black arrows denote surface spikes and epitope insertion points, where lower electron density is observed. Protein is black. Scale bar 5 nm.

### Structural analysis of H1 nanoparticles

The structures of the scaffold and H1-nanoparticle were found to be similar in size and shape, both appearing to be approximately 30 nm icosahedral particles (Fig. S6B vs. S6D). We studied the structure by cryo-EM to further characterize the H1-nanoparticle and to identify the location of helix-A epitopes. Cryo-EM micrographs indicate homogenous H1-nanoparticles with no observed aggregation (Fig. 2C). Reference-free class averages indicate that icosahedral symmetry of the nanoparticle scaffold is maintained (Fig. 2D). The 3D reconstruction confirmed the icosahedral symmetry (Fig. 2E) and demonstrates marked similarity to the scaffold structure (Fig. S8A-D). While the structure of the capsid is clearly resolved, the insertion site of helix-A lacks the strong density of the scaffold at the extruding spikes, indicating that the helix-A insertions are flexible or have multiple conformations (Fig. 2E, arrows). This is in stark comparison to the scaffold alone, which does not show any additional density above the capsid (Fig. S8C vs. S8D). The observed weak cryo-EM density at the insertion site is consistent with the homology modeling, which suggested flexible helix-A inserts and a relatively stable scaffold core (Fig. 2E, S5).

### Heat stability and immunogenicity of nanoparticle-presented epitopes

The strong density of the core of the H1-nanoparticle observed by cryo-EM suggested that the nanoparticle may be a stable and robust platform for epitope display. To test this, we probed for the impact of temperature fluctuations on nanoparticle stability and immunogenicity. Purified H1-nanoparticles aliquots were stored at 4°C, as would be done in a traditional cold chain. Additionally, aliquots were heated to temperatures of 40°C or 90°C for one hour and then all aliquots were equilibrated to room temperature. While there are some visual differences like levels of stain penetration (Fig. 3A vs 3B) and some aggregation (Fig. 3C), the images obtained by negative-stain electron microscopy illustrated that nanoparticles remained intact for all temperature probes (Fig. 3A, 3B, 3C).

**Fig. 3.**
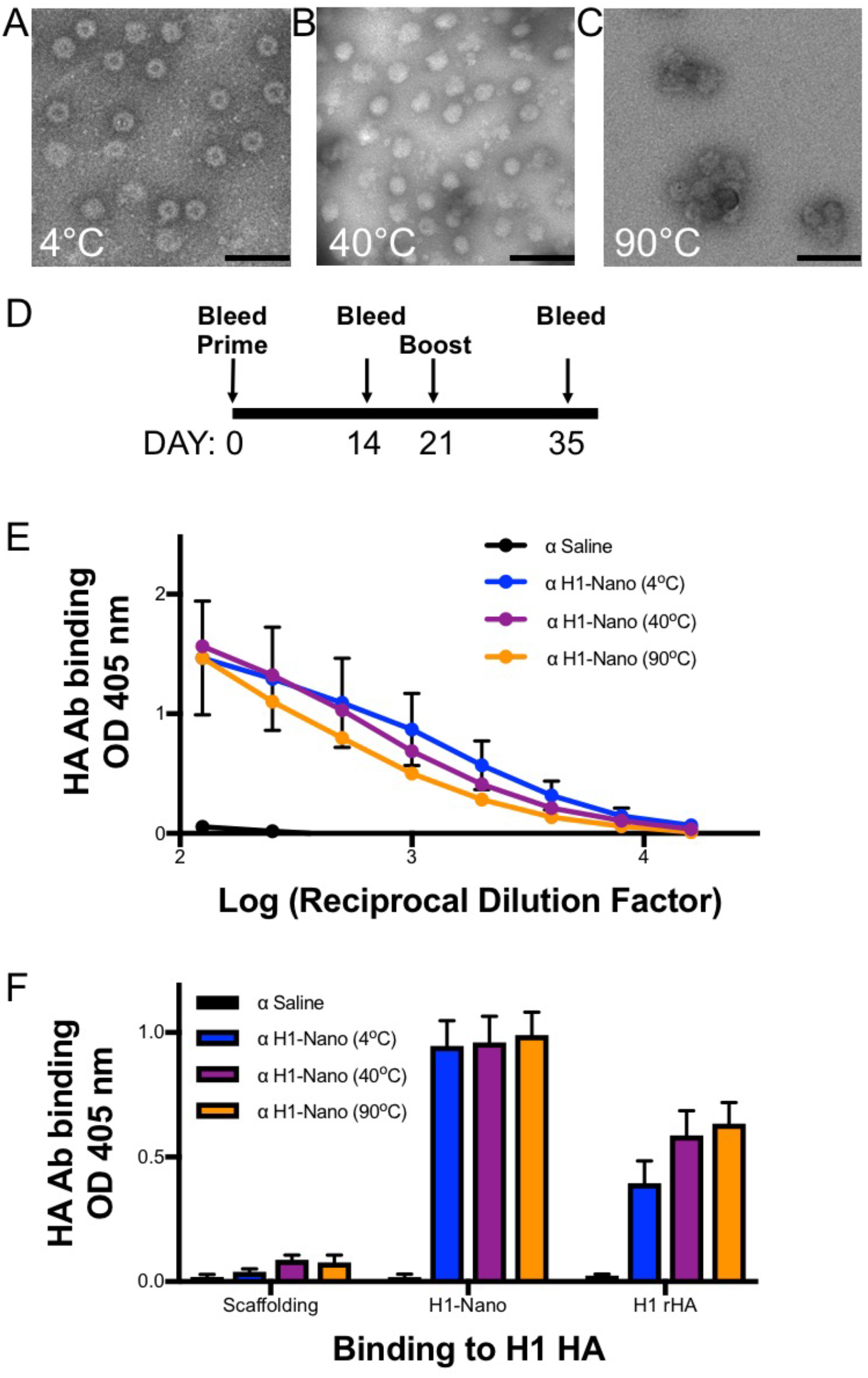
Probing the effect of temperature on H1 nanoparticle integrity and immunogenicity. (A, B, C) Negative-stained electron microscopy images of nanoparticles incubated at 4, 40, and 90°C. Particles were incubated for 60 min and then equilibrated to 25°C. Scale bars 100 nm. There are observed differences in stain penetration of the nanoparticles with a central cavity observed for Nano 4 (panel A) while there is less stain penetration and smoother looking particles for Nano 40 (panel B). Some aggregated particles appear in Nano 90 (panel C). (D) Schedule for immunization of mice with H1-nanoparticles with day 35 sera used in ELISA. (E) Comparison of sera reactivity to full-length recombinant H1 HA protein (H1 rHA) from mice immunized with H1-nanoparticles via ELISA with serially diluted sera. (F). Comparing the levels of reactivity of different sera for reactivity to scaffold, H1-nanoparticle (H1-Nano) and full-length recombinant H1 HA protein (H1 rHA). There were four groups of sera tested consisting of PBS (saline) and H1-nanoparticle exposed to three temperatures 4, 40, and 90°C (Nano 4, 40, 90). ELISAs for panels E and F are independent experiments.

A mouse model was used to assess the immunogenicity of purified nanoparticles with and without heating. Mice were primed (day 0) and boosted (day 21) with H1-nanoparticles that had been previously temperature treated (Fig. 3D). Mouse sera (day 35) following boost on day 21 was analyzed for reactivity with the following proteins by ELISA: scaffold, H1-nanoparticle, and recombinant H1 HA protein (rHA). A dilution series was used to identify the concentration of sera to be used for subsequent comparisons (Fig. 3E). The dilution series showed that compared to saline (black), sera from animals treated with nanoparticles kept at 4°C (blue) showed reactivity to H1 rHA (p = 0.0366), indicating that the nanoparticles are immunogenic. Sera from animals treated with nanoparticles that were heated and then equilibrated at room temperature before injection (purple and orange) showed similar reactivity to H1 rHA. For all subsequent ELISA experiments, sera was diluted 1:1000 in PBS. Dilution series and single dilutions of antigens in ELISA assays are separate independent experiments (Fig. 3E, 3F). Further analysis of the sera by ELISA showed that there was no significant difference in sera binding between the heated nanoparticles and those that maintained cold chain. For all three samples, antibodies were elicited not only to H1 rHA as would be needed for a successful vaccine candidate, but also to the H1-nanoparticle itself (Fig. 3F). However, there was only limited reactivity to the scaffold, likely because insertion of the helix-A epitope disrupted the immunodominant loop of the HBV capsid.

### Epitope identification of nanoparticle-elicited antibodies

To determine if antibodies elicited by the nanoparticle were specific to the stem of HA, sera from mice immunized with H1-nanoparticle kept at 4°C were analyzed against different H1 HA recombinant proteins by ELISA and western blot. ELISA analyses showed that sera had the same binding profile as a mouse monoclonal stem antibody (C179) when binding trimeric full-length HA (HA0) and the trimeric HA ectodomain which lacks the transmembrane region of HA (Fig. 4A). Like the monoclonal stem antibody, the nanoparticle sera did not react with recombinant HA1 head protein (rHA), which does not contain the HA stem region (Fig. 4A). Mice immunized with the H1-nanoparticle using the schedule described previously elicited antibodies that detected the H1-nanoparticle (~27 kDa) and H1 rHA (~70 kDa), but only weakly detected the scaffold protein (~20 kDa) (Fig 4B). This data aligns with the ELISA results for the pandemic H1-nanoparticle.

**Fig. 4.**
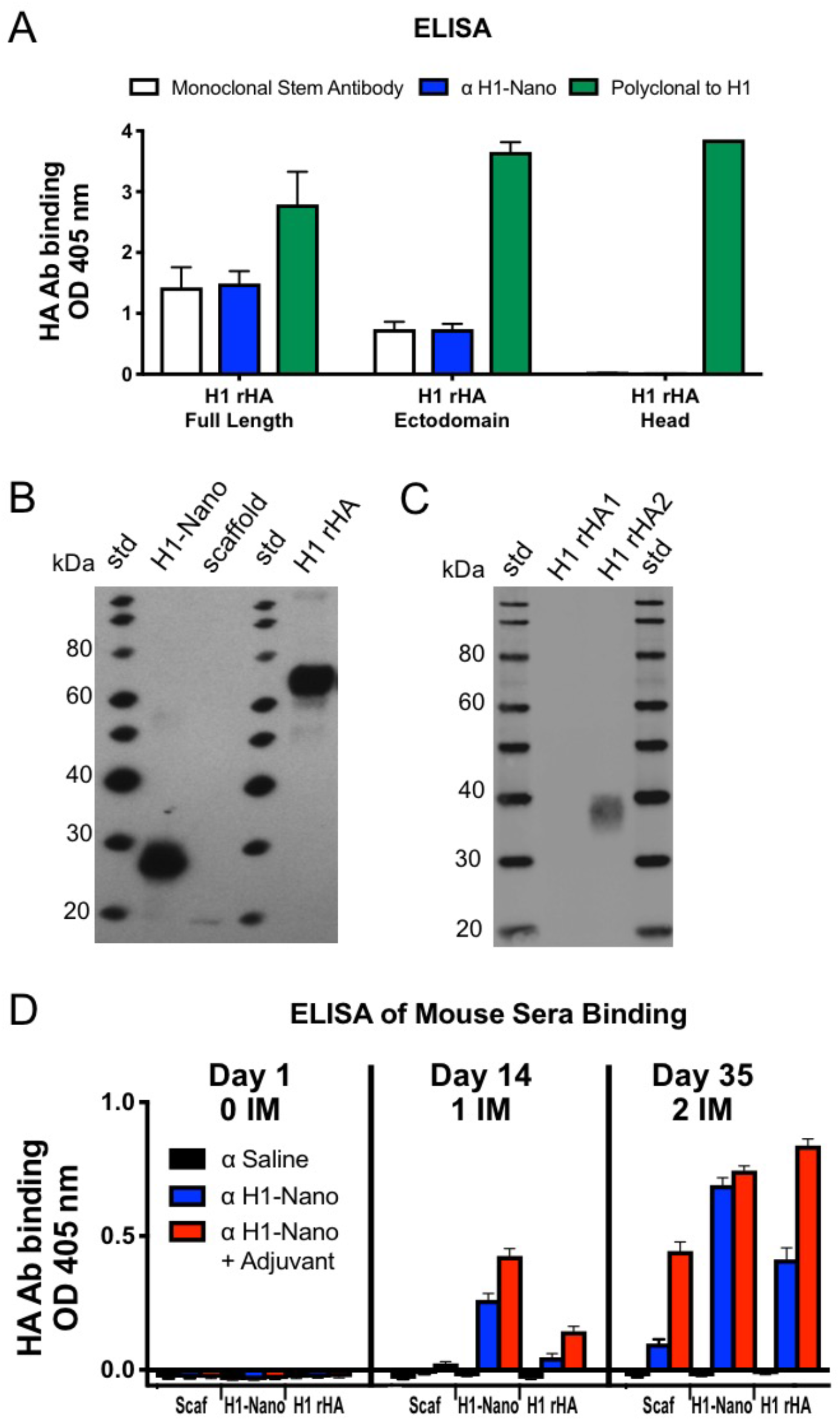
Immunogenicity of H1 nanoparticle and analysis of sera reactivity to different proteins. (A) ELISA binding analysis of day 35 sera from mice immunized with H1-nanoparticles binding to different recombinant H1 HA proteins: H1 rHA full-length, H1 rHA ectodomain and H1 rHA1 head domain (blue bars). Controls were a mouse monoclonal stem antibody (C179, white bars) and a rabbit polyclonal serum to H1 HA (green bars). (B) Reactivity analysis via western blot of sera from mice immunized with H1-nanoparticle alone to H1-nanoparticle (H1-Nano), scaffold, recombinant H1 HA (H1 rHA), and (C) to recombinant H1 rHA1 head and H1 rHA2 stem proteins. (D) Evaluation of serum antibody reactivity from immunized mice for days 0, 14, and 35. There were three groups of sera tested, which consisted of saline (black bar), H1-nanoparticle without adjuvant (blue bar), and H1-nanoparticle with adjuvant (red bar). Sera was tested for reactivity by ELISA to scaffold (Scaf), H1-nanoparticle (H1-Nano) and full-length recombinant H1 HA protein (H1 rHA). Recombinant proteins were from H1 HA of influenza (A/California/07/09) H1N1 and adjuvant was Sigma Adjuvant System (oil-in-water emulsion). H1 rHA stem ectodomain protein was based on a group 1 HA 1 stalk (stem) construct #4900, (Impagliazzo 2015).

To probe whether antibodies elicited by our nanoparticle platform could bind to the influenza stem, we used a group 1 stem (H1) construct of a headless stem trimer composed mostly of HA2 which has been previously used as a probe for stem antibodies in sera (29, 56). By western blot, the H1-nanoparticle sera did not detect the HA1 head domain, but did detect the recombinant HA2 stem construct (~40 kDa) (Fig. 4C). These results, combined with the results from the ELISA experiments, suggest that this nanoparticle platform displaying helix-A elicits antibodies that are reactive to a continuous epitope within the stem region of HA.

### Effect of adjuvant on immunogenicity

To determine the effect of adjuvant on the immunogenicity of the nanoparticles, we repeated the mouse immunization experiments using pandemic H1-nanoparticles in the presence and absence of the Sigma Adjuvant System. Statistical analysis was carried out for analysis of variance (ANOVA) using an F-test. The F statistic is reported (F=) with the first number in parenthesis being the degrees of freedom between groups and the second number the degrees of freedom within groups separated by a comma along with and the significance level (p).

We found that the adjuvant increased overall antibody levels (F (16, 510) = 137.9, p < 0.0001). However, there is no difference in the antibodies elicited to the nanoparticle itself with or without adjuvant after boosting (Fig. 4D, Day 35, H1-Nano). Nevertheless, both non-adjuvanted and adjuvanted H1-nanoparticle groups had at least a 4-fold increase in antibodies to H1 rHA after boosting, showing the importance of the prime-boost schedule (Fig. 4D, Day 14 vs. Day 35, H1 rHA,). The addition of adjuvant increased antibody binding to H1 rHA after one (F (4, 171) = 48.77, p < 0.0001) or two administrations of the nanoparticles (F (4, 168) = 51.65, p <0.0001).

### H1-nanoparticle protection from viral challenge

To determine if the nanoparticle-elicited immune response can protect animals from virus challenge, mice were immunized with the pandemic H1-nanoparticle with or without adjuvant and challenged with the H1N1 2009 pandemic virus. The immunization-challenge schedule was consistent with the immunogenicity studies from day 0 to 35. Additionally, mice were challenged at day 42 with 10x MLD_50_ (Mouse Lethal Dose) of pandemic H1N1 virus (A/California/07/2009) and a terminal bleed was performed at day 56 for the surviving mice (Fig. 5A). Mice immunized with non-adjuvanted H1-nanoparticle had a survival rate of 80% after challenge, while those animals immunized with adjuvanted H1-nanoparticle had a survival rate of 40% (Fig. 5B). Weight loss in both cases was reduced compared to the PBS control mice (Fig. 5C). Despite immunogenicity studies showing adjuvanted nanoparticles have more binding to H1 rHA protein (Fig. 4D, Day 35), these viral challenge results show that the adjuvanted nanoparticles do not protect mice from virus challenge better than non-adjuvanted nanoparticles.

**Fig. 5.**
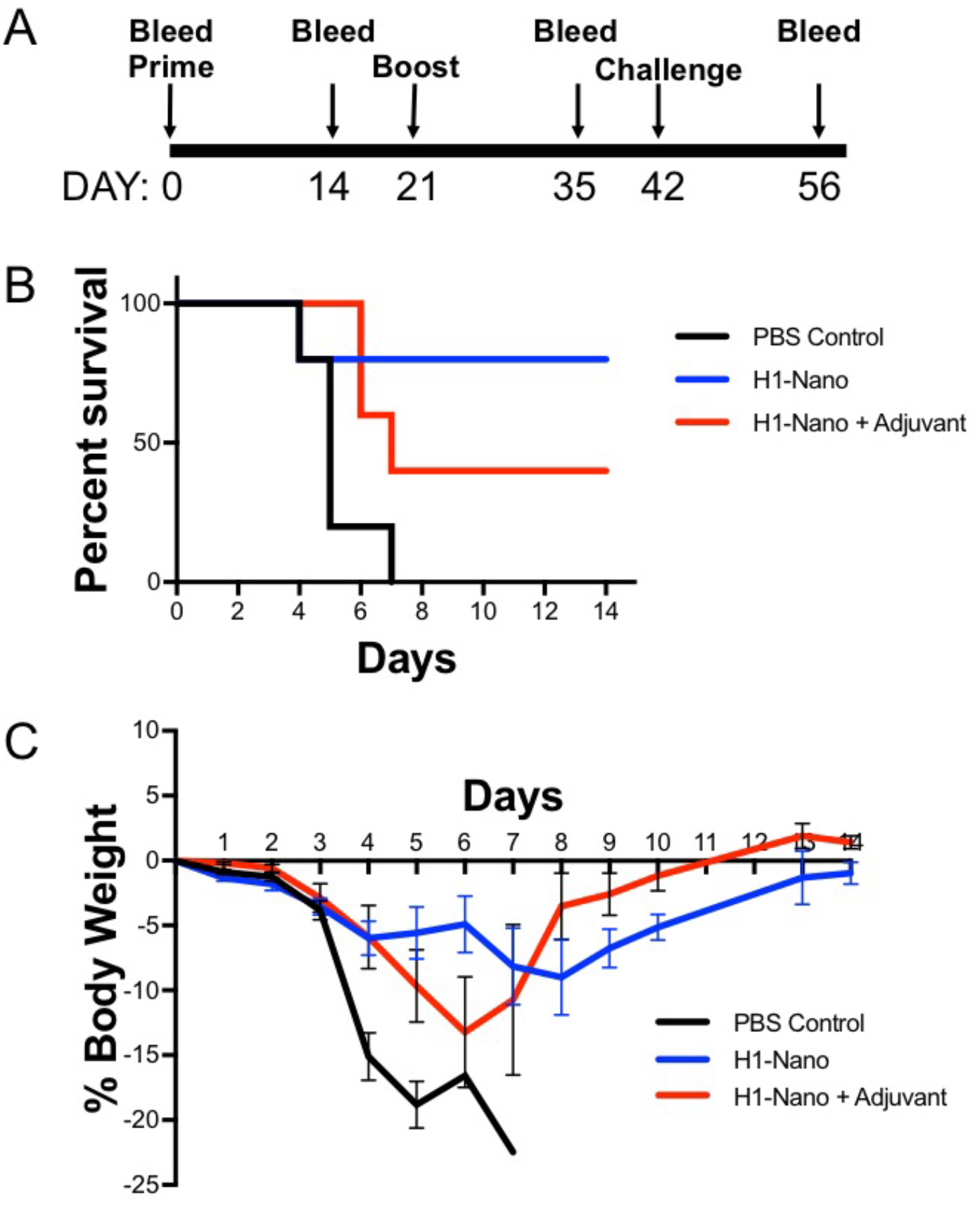
Assessment of H1 nanoparticle efficacy in mice. (A) Schedule for mouse immunization with H1-nanoparticle and challenge with H1N1 influenza virus. Groups of mice (N=5 per group) received one of three types of intramuscular injections: PBS, H1-nanoparticle without adjuvant or H1-nanoparticle with adjuvant on day 0 and 21. Adjuvant was Sigma Adjuvant System. Mice were challenged with 10x MLD_50_ (Mouse Lethal Dose) of H1N1 (A/California/07/2009) virus on day 42. (B) Survival curves for mice immunized with PBS (black), H1-nanoparticle (blue), or H1-nanoparticle with adjuvant (red) after challenging with virus. (C) Weight-loss curves for challenged mice that were immunized with PBS control (black), H1-nanoparticle (blue), or H1-nanoparticle with adjuvant (red).

To further characterize antibodies elicited in the challenge experiment, the sera was assessed for hemagglutination-inhibition (HAI) activity and microneutralization (MN) activity (Fig. S9). We observed negligible activity by HAI (Fig. S9B) or MN (Fig. S9C) from sera prior to virus challenge, suggesting that protective antibodies were not elicited to the head region and, consequently, were directed to HA stem epitopes. However, mice that survived challenge had HAI and MN titers at levels above background (>10 HAI, >20 MN) because the virus exhibits a head-dominant immune response (Fig. S9). These results, coupled with prior immunoassay analyses, indicate that the nanoparticles elicit antibodies to the stem region of HA and provide protection from influenza infection in a challenge model.

### Breadth of nanoparticle antibody response

Since the helix-A region of the stem is conserved between HA subtypes, we hypothesized that animals immunized with the H1-nanoparticle may have cross-reactive binding to other HA subtypes. We tested the H1-nanoparticle sera for binding to H7 rHA, a group 2 hemagglutinin that is found in the H7N9 virus and has pandemic potential. As a control for homosubtypic binding, we produced and purified H7-nanoparticles using the helix-A sequence from A/Anhui/01/2013 (H7N9) and immunized mice (Fig. S10). Mice were immunized following the previously described schedule (Fig. 3D).

Mice immunized with the pandemic H1-nanoparticles elicited antibodies capable of binding not only to H1 rHA (Fig. 4) but also to H7 rHA protein by ELISA compared to mice immunized with saline (p < 0.0001) (Fig. 6A), although H7 nanoparticle sera binding to H7 HA (p < 0.0001) was higher than H1-nanoparticle sera (Fig. 6A, pink vs blue line). H1-nanoparticle sera was also able to bind H7 rHA protein by western blot (Fig. 6B).These results indicate that antibodies elicited by influenza group 1 nanoparticles are able to cross-react with group 2 HAs.

**Fig. 6.**
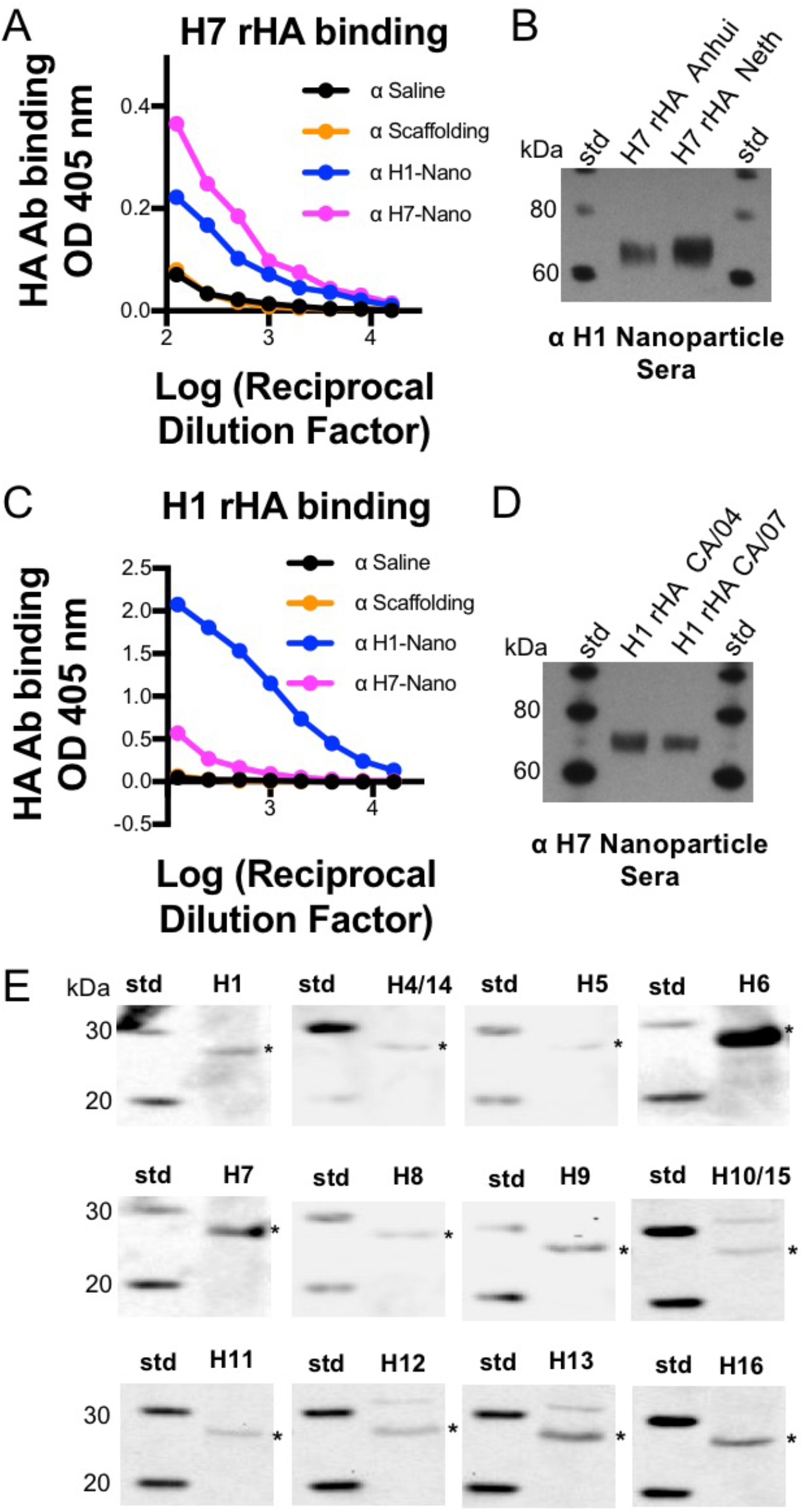
Probing sera for homosubtypic and heterosubtypic antibody reactivity to recombinant HA proteins. (A) Binding by ELISA of sera from mice immunized with different HA helix-A nanoparticles to recombinant H7 HA (A/Anhui/01/2013). (B) Western blot of sera from mice immunized with H1-nanoparticle (group 1 HA) to recombinant H7 HA proteins (group 2 HA) from H7N9 (H7 rHA A/Anhui/01/2013), and H7N7 (H7 rHA A/Netherlands/219/2003) viruses. (C) Binding by ELISA of sera from mice immunized with different HA helix-A nanoparticles to recombinant H1 HA (A/California/07/2009). (D) Western blot of sera from mice immunized with H7 nanoparticle (group 2 HA) to recombinant H1 HA proteins (group 1 HAs, H1N1 A/California/04/2009, A/California/07/2009). (E) Assessment by western blots of expression of designed helix-A HA-nanoparticle proteins from different HA subtypes from a nanoparticle library as indicated. Antibody 10E11 which binds an epitope tag in the scaffold was used as primary antibody. Each hemagglutinin subtype is denoted as H1, H4., etc., and asterisks denote detected bands. HA subtypes H4 and H14 have the same helix A as does HA subtypes H10 and H15 and they are represented by the same helix-A nanoparticle construct (H4/H14, H10/15).

To determine if influenza group 2 nanoparticles could elicit cross-reactive antibodies, we tested binding of sera from mice immunized with the H7-nanoparticles for binding to pandemic H1 rHA proteins. Compared to antibodies from mice immunized with saline, H7-nanoparticle sera antibodies were able to bind to H7 rHAs (Fig. S10), and interestingly also to H1 rHA. Although ELISA binding could be judged to be weaker for heterosubtypic HA binding than for homosubtypic HA binding, both ELISA (p < 0.0001) (Fig. 6C) and western blot (Fig. 6D) indicated H7-nanoparticle sera binding to H1 HA recombinant protein. These observations suggest that the helix-A nanoparticle platform can elicit homosubtypic and heterosubtypic antibodies for both group 1 and group 2 influenza subtypes.

Encouraged by these results, this design template was used to develop a library of nanoparticles that integrate helix-A epitopes from different HA subtypes (H1-H16) that includes avian and human viruses. We screened small-scale productions of constructs for all of these subtypes by western blot using primary antibody to an epitope tag region in the scaffold. Screening results indicated expression of nanoparticles for at least fourteen subtypes: H1, H4, H5, H6, H7, H8, H9, H10, H11, H12, H13, H14, H15, and H16 (Fig. 6E). Because HA subtypes H4 and H14 have the same helix A, as does HA subtypes H10 and H15, they are represented by the same helix-A nanoparticle (Fig. 6E, H4/H14, H10/15). Future studies will investigate the breadth of these nanoparticles for eliciting cross-protective antibodies, towards the generation of more effective seasonal, pandemic and a universal influenza vaccine.

## Discussion

In this work, we engineered novel chimeric nanoparticles using structure-guided design to combine the helix-A sequence from pandemic H1 HA with an HBV nanoparticle scaffold. We assessed feasibility of design and production, analyzed the structure and stability, and investigated the immunogenicity and protective capacity of the engineered nanoparticles. Our results indicate that nanoparticles displaying small non-trimeric HA2 stem epitopes (i.e. helix-A) can be expressed and purified, are heat-stable and are immunogenic for both group 1 and group 2 HA subtypes. Furthermore, this platform can be expanded to express helix-A nanoparticles for most HA subtypes. These results are valuable to vaccine development strategies that seek to universally increase protection from different influenza subtypes. Additionally, our results will aid in the development of vaccines with increased stability, having the potential to increase shelf-life.

### Epitope selection and copy number on nanoparticle scaffolds

An emerging strategy in vaccine development is to use symmetrical nanoparticles to display a viral antigen with multiple copies of the antigen displayed to the immune system (31, 32, 57-59). Repetitive antigen arrangements on nanoparticle surfaces is thought to facilitate B cell receptor co-aggregation, triggering, and activation (60). Studies have shown that multivalent viral proteins displayed on nanoparticles are more immunogenic than other platforms (31, 57, 58, 61, 62). Although the nanoparticle display of epitopes appears to be an emerging solution to invoke multivalent display, several factors must be considered, such as the identity, molecular size, density/spacing, and copy number of the epitope, as well as the size and type of nanoparticle scaffold. These parameters will affect epitope display and may alter the immune response. Our nanoparticle design allows for stem epitopes to have spacing that include 5 to 10 nm (Fig. 1G). This is important because 5-10 nm epitope spacings have been suggested to be the optimal distances to cross link B cell receptors (63).

When choosing a nanoparticle scaffold, it is important to minimize the risk of eliciting antibodies that could be cross-reactive to human antigens via vaccine-induced autoimmunity. Scaffolds of bacterial origin such as ferritin, encapsulin, and lumazine synthase maximize sequence divergence from similar proteins in humans to avoid this consequence (57, 58). We chose the naturally occurring HBV capsid as our nanoparticle scaffold, which is a viral capsid and thus not innately found in the human genome. For the helix-A nanoparticle sera very weak cross-reactive antibody to the HBV capsid scaffold was detected (Fig. 3F). This is probably due to disruption of the HBV capsid immunodominant site via HA helix-A insertion and there may be less cross-reactivity in persons with HBV capsid antibodies. Also, other non-human hepadnavirus capsids could be used as scaffolds if necessary because capsid sequences are divergent but share a similar structure (64). HBV forms icosahedral nanoparticles from 240 subunits of the capsid protein (65, 66), therefore we could integrate 480 copies of helix-A on each nanoparticle (Fig. 1G). This represents a significant increase in the HA stem epitope copy number as compared to other nanoparticle influenza vaccine platforms using trimers (35).

Our designed helix-A nanoparticles used the HBV viral capsid as a scaffold. This differs for other studies that use computational methods to design sequences for scaffolds that form nanoparticles with desired properties (67-69), as has been done for respiratory syncytial virus (RSV) (70, 71) (62). While these strategies have shown promise, designing novel nanoparticle platforms is time-intensive and requires much more validation than naturally-occurring scaffolds. The strengths and weaknesses of naturally occurring scaffolds (e.g. viral capsids) and scaffolds designed de novo will require more investigation. For any specific design, the scaffold and epitope pairing will depend on parameters such as the particular epitope target, viral system, desired immune response, and stability requirements.

### Stability of influenza vaccines

Current influenza vaccines require refrigeration and vaccine potency can be lost upon improper storage and failure of the cold chain (13). Furthermore, the stability of recombinant influenza HA subunit vaccines correlates with the vaccine strain. For example, after vaccines lots were released for the 2009 pandemic H1N1 virus, there was an unexpected decrease in HA potency and the reported shelf life was shortened. Biochemical analysis revealed that this H1 HA strain was less heat-stable, with variation in glycosylation and susceptibility to protease activity (72). High-stability antigens may be valuable in eliminating concerns about the cold-chain and improving shelf-life for distribution of seasonal, pandemic, and universal influenza vaccines. The nanoparticle platform described here appears to have robust stability properties, displaying short term temperature stability to 90°C while maintaining immunogenicity of the helix-A epitope for the 2009 pandemic H1N1 construct (Fig 3). Thus, our approach has potential advantages over commercially available vaccines.

A number of studies have suggested approaches to improve vaccine stability. These studies have tested the efficacy of microneedle patches to administer vaccines over a longer time period, as well as chemical cross-linking and additives to improve storage stability (73-76). While these studies have shown an increase in stability, the preservation of immunogenicity of epitopes must also be assessed. In our work, we demonstrate stability through integration of an immunogenic epitope with a stable nanoparticle scaffold. This nanoparticle maintained immunogenicity of the stem epitope even after heating, suggesting the plasticity of this HA stem nanoparticle design. An important future consideration for our work and other stability studies is how long the nanoparticle vaccines remain stable and whether these vaccines maintain immunogenicity in humans.

### Expanding stem nanoparticle design and expression to more HA subtypes

Our pandemic H1 helix-A stem nanoparticle is immunogenic and protects from challenge in mice. Only two immunizations were required and an oil-in-water emulsion adjuvant appears not to increase protection. This will require further studies to understand, since this differs from studies of trimeric antigens, which require adjuvant and, in some cases, require three immunizations (29, 35, 49-52). This design concept can be applied more broadly to other influenza A HA subtypes, as we have demonstrated expression for the majority of influenza A HA subtypes (Fig. 6E). At present it appears that trimeric stem immunogens have been reported for only H1, H3, H5, and H7 (29, 35, 49-52).

It may be challenging to produce traditional trimeric stem immunogens for all subtypes that successfully fold into an immunogenic epitope. Previous studies have shown that stabilizing mutations and trimerization motifs may be required to maintain biological relevance of a trimeric stem epitope (29, 35, 49-52). Consequently, numerous constructs may need to be screened for expression, stability, and structure (29, 35). Our approach did not require amino acid changes within the stem region epitope, helix-A. Additionally, we were able to identify helix-A sequences through computational methods to develop a library of nanoparticles that cover the majority of HA subtypes (Fig. S3). Targeting additional HA subtypes may open opportunities to improve immunogenicity and broaden protective immune responses to different strains, subtypes, and types of influenza virus.

Another way to target additional subtypes of HA is through cross-protection, which is important both for safeguarding against antigenic drift and in developing a universal influenza vaccine. In this study, we showed that the pandemic H1 helix-A stem nanoparticle elicited antibodies that could bind to phylogenetically distinct group 1 (H1) and group 2 (H7) HA proteins (Fig. 6A-6D). We have expanded design of our nanoparticle platform to all HA subtypes, allowing expanded future studies on the ability of the helix-A nanoparticle platform to elicit antibodies that target more HA subtypes. In future studies, it will be interesting to test whether the expanded library of helix-A HA stem nanoparticles elicit homosubtypic and heterosubtypic cross-reactive antibodies to different HA subtypes. Further, it will be important to test whether the cross-reactive antibody responses generated offer broad protection from viral challenge. This will be important for optimizing broad immune responses to the conserved stem region of antigenically different HAs for universal influenza vaccine development. One notion is that combinations of nanoparticles displaying conserved stem epitopes of different HA subtypes given together in a vaccine cocktail may further enhance the breadth of protection for a more universal vaccine.

In conclusion, our results indicate that helix-A sequences are conserved and can be displayed as antigenic epitopes on a nanoparticle scaffold. Using pandemic H1-nanoparticles as a proof of concept, we showed that the helix-A epitope is immunogenic and that the nanoparticles were protective in an H1N1 challenge model. Furthermore, the elicited antibodies exhibited homosubtypic and heterosubtypic binding between group 1 (H1) and group 2 (H7) HA proteins. We also established an expression library of helix-A nanoparticles for the majority of HA subtypes. Due to the broad array of influenza subtypes that can be targeted and their robust properties, our results suggest that helix-A nanoparticle immunogens should be further explored to aid in the development of more efficacious and broadly protective influenza vaccines.

## Materials and Methods

### Informatics and nanoparticle DNA construct designs

In order to better determine the antigenic variation in the helix-A stem region (A-helix), a time snapshot of the amino acid sequences for all fludb.org influenza hemagglutinin (HA) sequences was categorized. The database was divided into individual databases for H1 – H16, the amino acid sequence distributions of the helix A regions were compiled for each HA subtype. The consensus sequence was determined per HA subtype. Further, the identity and occurrence of each variation from the consensus sequence was classified. Individual amino acid differences and total sequence identity variation between H1 – H16 consensus sequences was determined using a matrix to compare sequence identified between the helix-A consensus sequences (Fig. S2). Sequence identities were performed with EMOBSS-lite (formerly GCG-lite). Helix-A nanoparticles were designed by making chimeric protein sequences by inserting two copies of helix A from HA stem regions from 27 selected sequences representing H1-H16 subtypes into the immunogenic loop region the human hepatitis B virus capsid. The HBV sequence reference was YP_355335 (30-212) with the precore region removed. The capsid was used as a scaffold. Linkers (G-G-G) flanked the helix-A sequences. The chimeric protein DNA sequences were then codon optimized for E. Coli expression and was then synthesized by Life Technologies into a pMA-T vector backbone. Plasmid designations were based on the HA sequence. For example, pMA-T-H01A contains DNA for an H1-nanoparticle for the helix A stem region from H1 HA A/California/07/2009 virus while pMA-T-H07B is for the helix A stem region of H7 HA from H7N9 A/Anhui/01/2013 virus.

### Molecular models of designed nanoparticle generated with RosettaCM

RosettaCM, a comparative modeling protocol within the Rosetta software package (77) was used to predict possible structures of a monomer of the H1-nanoparticle construct. The program samples multiple templates and uses secondary structure segments that may be recombined between templates to generate numerous possible models that can be evaluated based on a score related to its free energy. In this case, secondary structure segments were generated using the Robetta structure prediction server. Additionally, templates of the HBV capsid protein (PDB:1QGT) and H1 hemagglutinin sequence (PDB:3LZG) were used to generate the models. Template matching was performed such that the HBV sequence was aligned to the scaffold core only and each insert was aligned to an individual H1 HA template. 10,000 predicted models were generated and the 5% with the lowest free energy were selected for further evaluation. These were then analyzed for their ability to multimerize without significant clashes with neighboring monomers, based on aligning the scaffold core coordinates to those from the HBV icosahedral capsid structure. Only those that did not have significant clashes were selected.

### Nanoparticle expression and purification

For production and purification of helix-A stem nanoparticle (e.g. H1-nanoparticle, H7-nanoparticle), the H1 and H7 constructs (pMA-T-H01A, pMA-T-H07B) were subcloned into pET21b expression vectors to create (pET21b-H01A, pET21b-H07B). These plasmids were transfected into Rossetta 2 cells and this system was used for expression according to manufacturer’s instructions (EMD Millipore) for expression. The E. coli Rossetta cells were grown in 2xYT liquid media at 37°C until the OD600 was 0.8. Cells were then induced with 1 mM IPTG and expressed overnight at 25°C. Cells were then pelleted at 3300x*g* for 30 min and resuspended in ice-cold PBS and then sonicated to release the crude nanoparticle. Cellular debris was removed by pelleting via centrifugation at 16,300x*g* for 10 min. The crude supernatant containing nanoparticle was then precipitated with ammonium sulfate at 45% saturation via incubation overnight at 4°C, and the precipitant was resuspended in PBS. This nanoparticle preparation was then layered on a gradient with 30% - 50% - 70% sucrose steps and centrifuged at 34,000 rpm using a SW55 Ti rotor for 2.5 hours. Sucrose fractions were collected, and the location of the nanoparticles were identified by SDS-PAGE. Fractions containing the nanoparticles were dialyzed overnight in PBS and then purified in three tandem CsCl gradients centrifuged in an SW55 Ti rotor at 38,000x rpm for 24 hours. The gradient band containing the nanoparticles were collected and dialyzed in PBS overnight. HBV capsid particles were purified in a similar fashion as H1 and H7 nanoparticles. Endotoxin was removed from the samples by adding 2.5% Triton X-114, mixing at 4°C for 1 hr, incubating at 37°C for 10 min, centrifuging at 16,500x*g* for 10 min at 25°C, and collecting the upper aqueous phase. Three cycles of Triton X-114 endotoxin removal were performed per purification. Endotoxin removal was confirmed by using the LAL chromogenic Endotoxin Quantation Kit (Thermo Fisher Scientific, Waltham, MA) according to the manufacturer’s instructions.

### SDS-PAGE

Nanoparticle purity and protein composition was analyzed biochemically by SDS-PAGE. Samples were analyzed under reducing conditions with the reducing agent dithiothreitol (DTT) at final concentration of 100 mM and heated at 95°C for 10 minutes. Gels were stained overnight with coomassie blue stain (SimplyBlue SafeStain, Invitrogen, Carlsbad, CA) for protein band visualization. Gels were scanned and digitized into images with a gel documentation system (Enduro GDS, Labnet International, Edison, NJ).

### Electron microscopy

Negative-staining electron microscopy of nanoparticles was similar to that reported previously for other samples (78), with the exception that the samples were stained with 1.5 % phosphotungstic acid. Images were collected on a Tecnai-12 electron microscope with LaB6 filament operating at 100 kV (FEI, Hillsboro, OR) at a nominal magnification of 52,000x. Images were recorded on a 4k x 4k OneView camera (Gatan, Pleasaton, CA).

For cryo-electron microcopy, data for the H1-nanoparticle and HBV capsid (scaffold) were collected under the same conditions, except where noted. 3.5 μL of unstained nanoparticles were applied to glow discharged 200 mesh R2/2 Quantifoil Cu grids (Quantifoil, Großlöbichau, Germany) and plunge frozen using a Vitrobot Mark IV plunger (FEI, Hillsboro, OR). Samples were imaged under cryo conditions at 300 kV on a Titan Krios electron microscope (FEI, Hillsboro, OR). Images were collected using EPU software (FEI, Hillsboro, OR) on a Falcon 2 direct electron detector (FEI, Hillsboro, OR) using a 1.37967 Å pixel size (nominal magnification 59,000x). Images were collected with a defocus range of −0.75 to −3.25 μm and the electron dose was approximately 22 e−/Å2. For HBV capsid 3,065 images were collected, and 2,858 images were collected for the H1-nanoparticle.

### 3-D reconstructions

Data for HBV capsid (scaffold) and the H1-nanoparticle were reconstructed using the same parameters, except where noted. Movie-mode images were aligned and averaged using MotionCor2. CTFFIND4 (79) was used to correct images for the contrast-transfer function. For HBV capsid, 311,614 particles were picked and the 2D classification function in Relion 2 was used to select 102,643 particles for reconstruction. For the H1-nanoparticle, 83,621 particles were chosen and 2D classification was used to select 21,485 particles for reconstruction. The Auto-refine function of Relion 2 (80) was used to reconstruct HBV capsids and H1-nanoparticles, using spherical internal and external masks to isolate capsid density. Postprocessing in Relion 2 yielded a reconstructed 3D density map of 4.8 Å and 10 Å for HBV capsid and the H1-nanoparticle, respectively.

### Heat treatment

Nanoparticles were heat treated for stability and immunogenicity assessments. Aliquots of purified nanoparticles in PBS were maintained at 4°C, incubated at 40ºC for one hour, or incubated at 90°C for one hour. Each of the three aliquots was then equilibrated to room temperature. One fraction of each aliquot was then used for negative-stain electron microscopy analysis and another fraction used for immunogenicity studies.

### ELISAs

Antigen was applied to 96-well plates and incubated overnight at 4°C (1.25 μg/mL), following PBS washes and blocking (1% Omniblok, AmericanBio, Inc., Natick, MA and 0.1% Tween 20 in PBS), primary antibody was incubated for 2 hr at room temperature (serum samples were diluted 1:1000). Plates were washed and placed at 37°C for 1 hr with the addition of an HRP conjugated secondary antibody (goat anti-mouse IgG (H+L), Thermo Fisher Scientific, Waltham, MA). Colorimetric detection occurred for 15 min at room temperature and absorbance was read at 405 nm (1-Step ABTS, Thermo Fisher Scientific, Waltham, MA). Samples were run in quadruplicate. Statistical calculations were carried out in with the software Prism (GraphPad, La Jolla, CA).

### Western Blots

Purified recombinant full-length HA proteins were from Protein Sciences Corporation, Meriden, CT and HA H1 ectodomains were from the International Reagent Resource. Headless trimeric H1 stem protein was kindly provided by Jeffrey Taubenberger (56). HA proteins along with designed nanoparticles and HBV capsid (scaffold) preparations were denatured and heated prior to loading (2 μg) on an polyacrylamide gels. Samples were transferred to 0.2 μm nitrocellulose membranes and blocked for 1 hr (1% Omniblok, AmericanBio, Inc., Natick, MA and 0.1% Tween20 in TBS). Membranes were incubated overnight with serum (20 μL in 20 mL blocking buffer) elicited from the nanoparticle immunizations, or with primary epitope-tag antibody to the capsid scaffold (mAb 10E11) when screening for expression of designed nanoparticle constructions. Blots were washed and probed with either a HRP conjugated secondary antibody (goat anti-mouse IgG (H+L), Thermo Fisher Scientific, Waltham, MA) or fluorescent-labeled secondary antibody (goat anti-mouse IgG). Blots were then visualized using SuperSignal West Pico chemiluminescent substrate (Thermo Fisher Scientific, Waltham, MA) to expose film and finally digitally scanned into images or imaged with an Azure c600 imager.

### Immunogenicity

All animal experiments were performed under protocols approved by the Animal Care and Use Committee (ACUC) at the National Institute of Allergy and Infectious Diseases (NIAID). A combination of 30 female and 30 male BALB/c mice (Charles River Laboratories, Maryland), aged 8-10 weeks were randomly assigned to antigen groups and underwent two immunizations (Day 0 and Day 21) and tail bleeds (Day 0 and Day 14) prior to a terminal bleed on Day 35. Mice were injected intramuscularly with 50 μL of antigen at a concentration of 0.5 mg/mL for a final experimental dose of 100 μg/mouse. 30 male mice were used for the heat-treatment immunogenicity studies following the same regimen. The H7-nanoparticle immunogenicity experiments were performed on 5 female BALB/c mice. No gender differences were observed.

### Challenge

Experiments involving challenges were conducted using protocols approval by NIAID ACUC. 15 female BALB/c mice, aged 8-10 weeks, underwent the same inoculation protocol as the immunogenicity experiments’ (injections Day 0 and Day 21). On Day 35 mice were anesthetized and intranasal challenged with 10x LD_50_ (i.e. MLD_50_ (Mouse Lethal Dose)) of influenza H1N1 (A/California/07/2009). Mice were observed twice daily for survival criteria (humane endpoints observed following a 25% decrease from initial body weight) until Day 56, when all surviving mice were humanely euthanized. Differences in survival rates were compared using a Kaplan-Meier survival analysis (Graph Pad Prism, La Jolla, CA).

### Hemagglutination inhibition (HAI) assay

The sera and plasma were diluted 1:4 in RDE (RDE II, “Seiken”, receptor-destroying enzyme, cat. no. UCC-340-122, Accurate Chemical) and placed in a 37°C water bath overnight (18-20 hr). Sera were heat-inactivated at 56°C for 40 min and adsorbed on turkey red blood cells (TRBC) for 30 minutes, leading to a final serum dilution of 1:10. Turkey red blood cells (TRBC) were prepared by mixing 0.5% TRBC in PBS. For each antigen, 50 μl PBS/0.5% BSA per well was added to all wells of 4 adjacent columns of a 96-well V-bottom plate. 50 μl virus stock was added to the first well of the 2 first columns, mixed, and serially diluted down the columns using 50 μl. Fifty (50) μl was discarded from the last well. 50 μl TRBC was added, the plate was agitated, and the hemagglutination pattern read after 30 min incubation at room temperature. The HA value was established as the greatest dilution resulting in complete hemagglutination. Viral antigens were diluted in PBS/0.5% BSA to contain 4 hemagglutinating units (HAU) in 25 μl and the HA value verified as follows: 4 wells of a 96-well plate will be filled with 50 μl PBS/BSA solution. The top well was filled with an additional 50 μl of diluted virus solution (4 HAU/25 μl) and titrated to the last well in two-fold dilutions. 50 μl TRBC was added, the plate agitated, and the HAU read after 30 min incubation at room temperature. Each well of a 96-well V bottom assay plate was filled with 25 μl PBS/BSA. Prepared sera or plasma was added across the top row and diluted down the columns in two-fold dilutions. Each sample was tested in duplicate. 25 μl of virus was added to each well except the last column. The plates were agitated and incubated for 30 min at room temperature. 50 μl TRBC was added to each well followed by a 30-min incubation, after which the hemagglutination patterns were read by tilting the plates at a slight angle.

### Microneutralization (MN) assay

The virus stock was titrated by adding 0.5 log dilutions of the virus stock to a 96 well plate containing MDCK cells. The titer was calculated using the Reed-Muench formula. The plate was washed three times with wash buffer (PBS/0.1% Tween-20) and the primary antibody (anti-NP mouse monoclonal Ab, Millipore, cat. MAB2851) added in blocking buffer. The plate was incubated for 1 hr at room temperature, washed three times in wash buffer and blocking buffer (PBS/1%BSA/0.1% Tween-20) was added. The plate was incubated for 10 min at room temperature, washed three times and the secondary antibody (goat anti-mouse IgG HRP, KPL, cat. no. 474-1802) was added in 100 μl/well. The plate was incubated for 1 hr at room temperature, washed three times with wash buffer, blocked in 200 μl blocking buffer as previously, and washed three times with wash buffer. The TMB substrate was added (100 μl/well) and incubated for 5-10 min at room temperature. The reaction was stopped with 100 μl of TMB stop solution (100μl/well) and the plate was read at 450 nm.

### Test Expression for H1-H16 HA from the nanoparticle library

Transformed E. coli cells containing plasmid expression vectors for 16 nanoparticle constructs representing H1-H16 HA subtypes were seeded into 4 mL of 2XYT liquid medium and induced with 1 mM IPTG to express overnight. Cells were subsequently isolated from medium via centrifugation (5000 rcf for 10 min) and lysed with B-PER (Thermo Fisher Scientific, Waltham, MA) for 15 min at room temperature; the content of the cell lysates were separated by centrifugation (16000 rcf for 7 min). Nanoparticle production was confirmed by western blot as described above using an epitope tag antibody (10E11) to the scaffold (capsid) region (aa. 2-10) that is in all the constructs.

## Acknowledgements

This work was supported by the Intramural Research Program of the National Institute of Allergy and Infectious Diseases. We thank Michael Conlon for help with protein purification, Jeffery Taubenberger and Jae-keun Park for kindly providing the headless trimeric stem protein and the International Reagent Resource for HA proteins. We thank Vinod Nair and Elizabeth Fischer for help in cryo-EM data collection. This work utilized the computational resources of the NIH HPC Biowulf cluster (http://hpc.nih.gov).

