## supplemental figures and captions for "Designed Nanoparticles Elicit Cross-Reactive Antibody Responses To Conserved Influenza Virus Hemagglutinin Stem Epitopes"

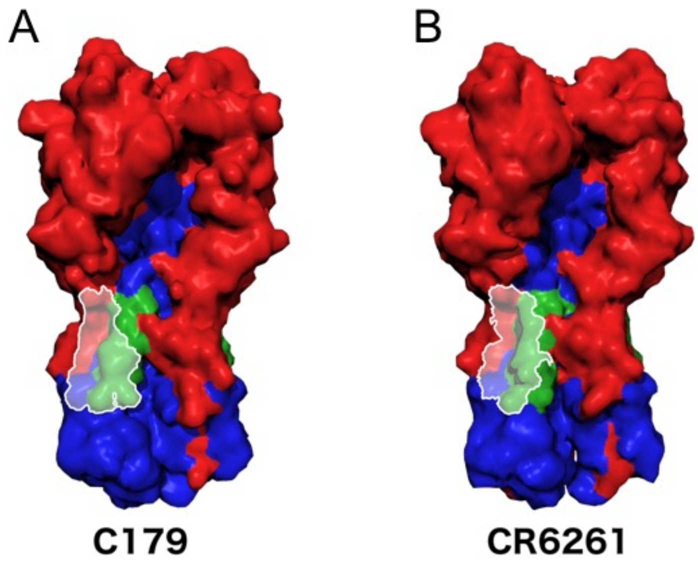


Fig. S1

Matrix of percent sequence identity between H1 to H16 hemagglutinin (HA) based on consensus sequences of Helix A of each HA subtype

| | H1 | H2 | H3 | H4 | H5 | H6 | H7 | H8 | H9 | H10 | H11 | H12 | H13 | H14 | H15 | H16 |
|---|---|---|---|---|---|---|---|---|---|---|---|---|---|---|---|---|
| H1 | | 86 | 64 | 68 | 73 | 77 | 68 | 68 | 59 | 68 | 55 | 59 | 59 | 68 | 68 | 59 |
| H2 | | | 55 | 59 | 86 | 86 | 64 | 64 | 64 | 64 | 68 | 64 | 73 | 59 | 64 | 68 |
| H3 | | | | 95 | 50 | 55 | 82 | 55 | 50 | 86 | 50 | 55 | 59 | 95 | 86 | 50 |
| H4 | | | | | 50 | 55 | 86 | 55 | 50 | 91 | 50 | 55 | 59 | 100 | 91 | 55 |
| H5 | | | | | | 91 | 55 | 68 | 68 | 55 | 73 | 64 | 77 | 50 | 55 | 64 |
| H6 | | | | | | | 59 | 73 | 77 | 59 | 73 | 68 | 77 | 55 | 59 | 64 |
| H7 | | | | | | | | 59 | 55 | 95 | 55 | 55 | 64 | 86 | 95 | 59 |
| H8 | | | | | | | | | 82 | 59 | 73 | 59 | 68 | 55 | 59 | 59 |
| H9 | | | | | | | | | | 55 | 82 | 68 | 73 | 50 | 55 | 64 |
| H10 | | | | | | | | | | | 55 | 55 | 64 | 91 | 100 | 59 |
| H11 | | | | | | | | | | | | 55 | 82 | 50 | 55 | 64 |
| H12 | | | | | | | | | | | | | 64 | 55 | 55 | 55 |
| H13 | | | | | | | | | | | | | | 59 | 64 | 82 |
| H14 | | | | | | | | | | | | | | | 91 | 55 |
| H15 | | | | | | | | | | | | | | | | 59 |
| H16 | | | | | | | | | | | | | | | | |

N=50,428 HA sequences (H1 through H16) from the influenza sequence database were analyzed. Identities ranged from 50% to 100%. Each HA subtype is represented by its consensus sequence.

Fig. S2

A

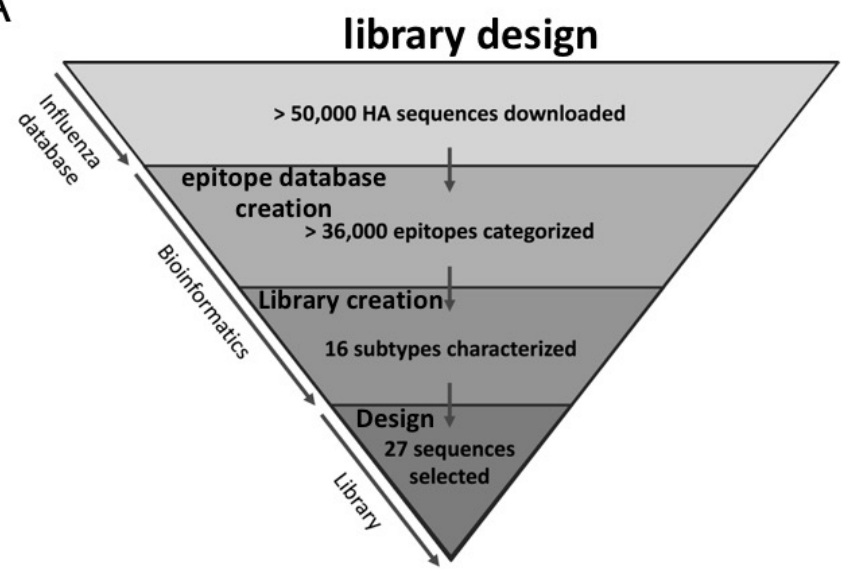


B

### Identity

| Percent identities of library to database sequences | Corresponding coverage of influenza database epitopes |
|---|---|
| 100% | 62.50% |
| 95% | 28.60% |
| 91% | 5.10% |
| 86% | 2.50% |
| 82% | 0.90% |
| 77% | 0.40% |

Fig. S3

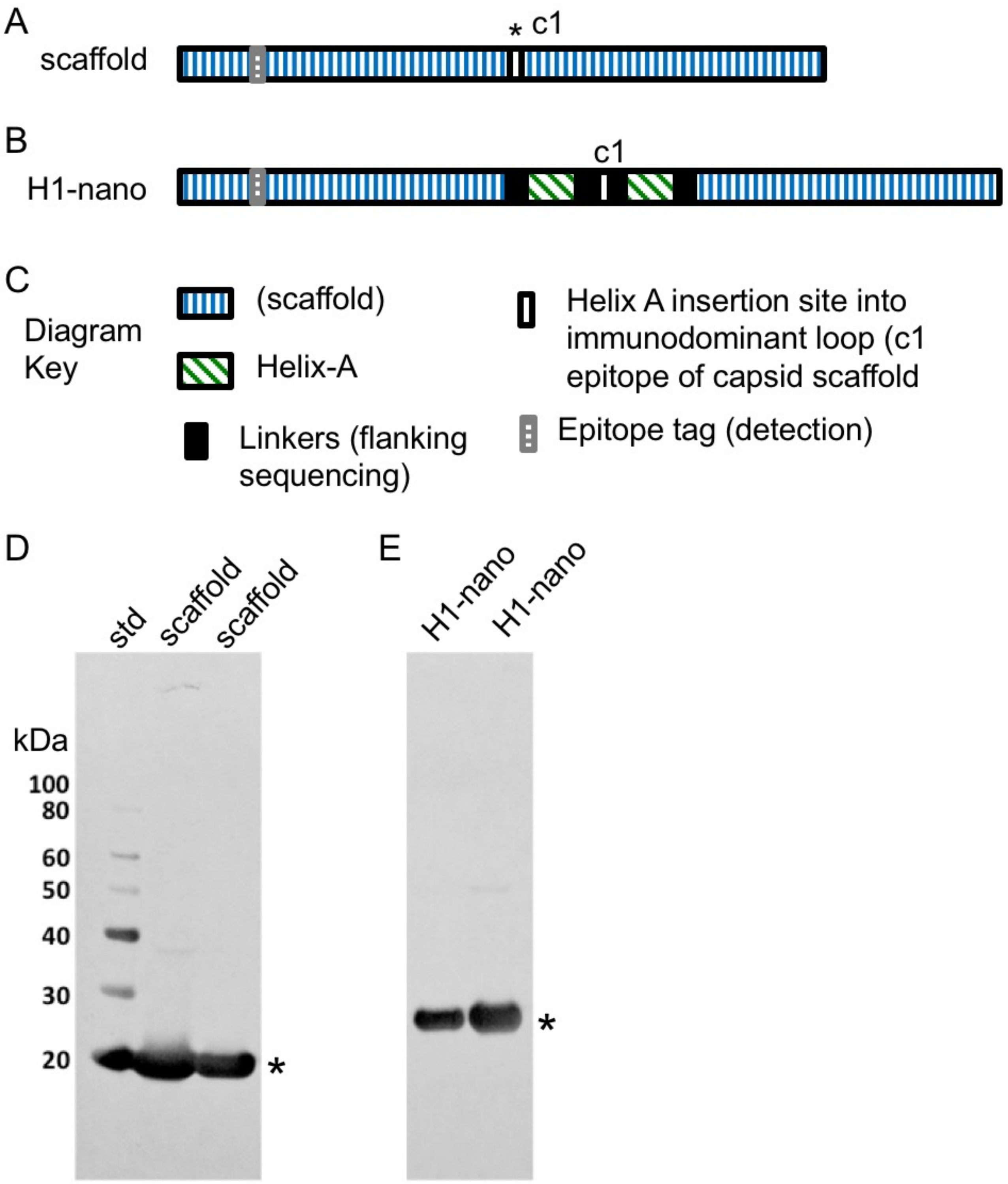
A
scaffold
* c1
B
H1-nano
c1
C
Diagram Key
(scaffold)
Helix-A
Linkers (flanking sequencing)
Helix A insertion site into immunodominant loop (c1 epitope of capsid scaffold
Epitope tag (detection)
D
std
scaffold
scaffold
kDa
100
80
60
50
40
30
20
*
E
H1-nano
H1-nano
*

Fig. S4

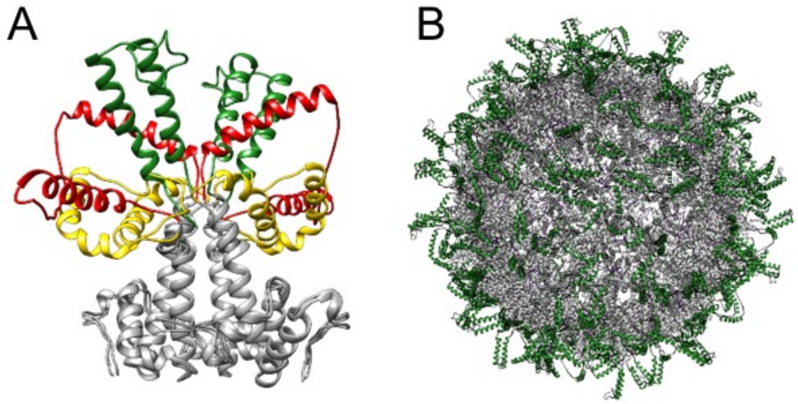


Fig. S5

Scaffold: Purification and particle formation

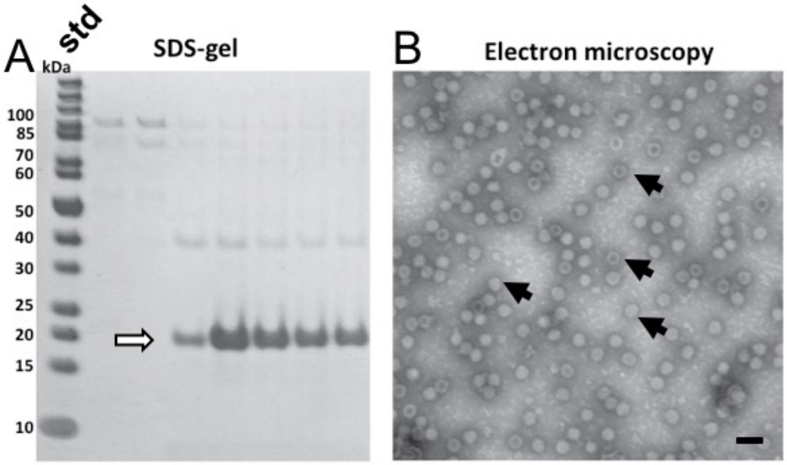


H1-nanoparticle: Purification and particle formation

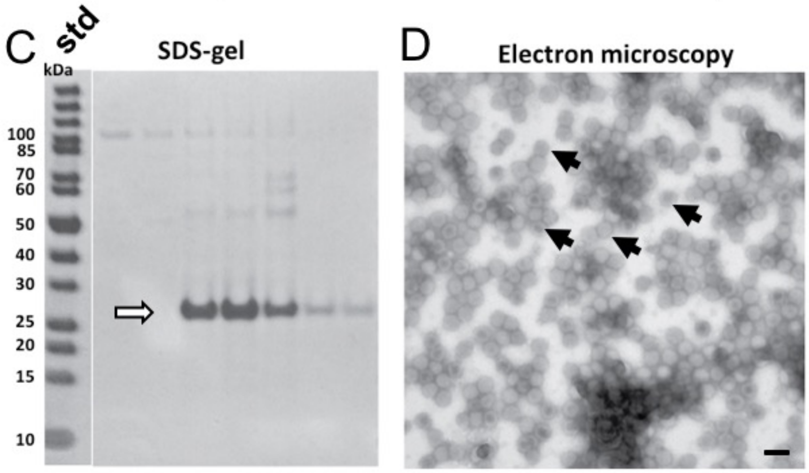


Fig. S6

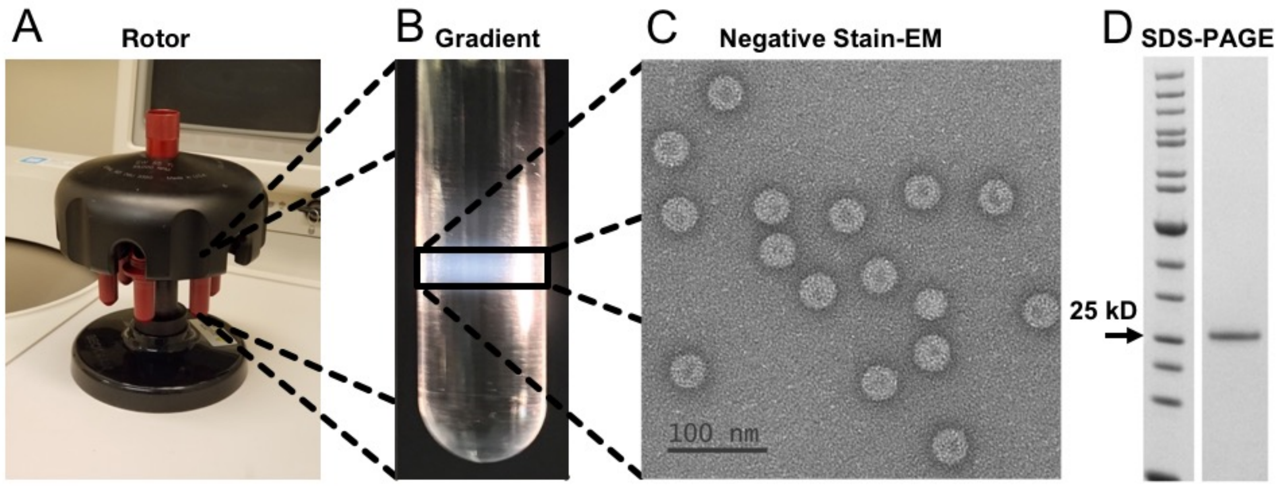


Fig. S7

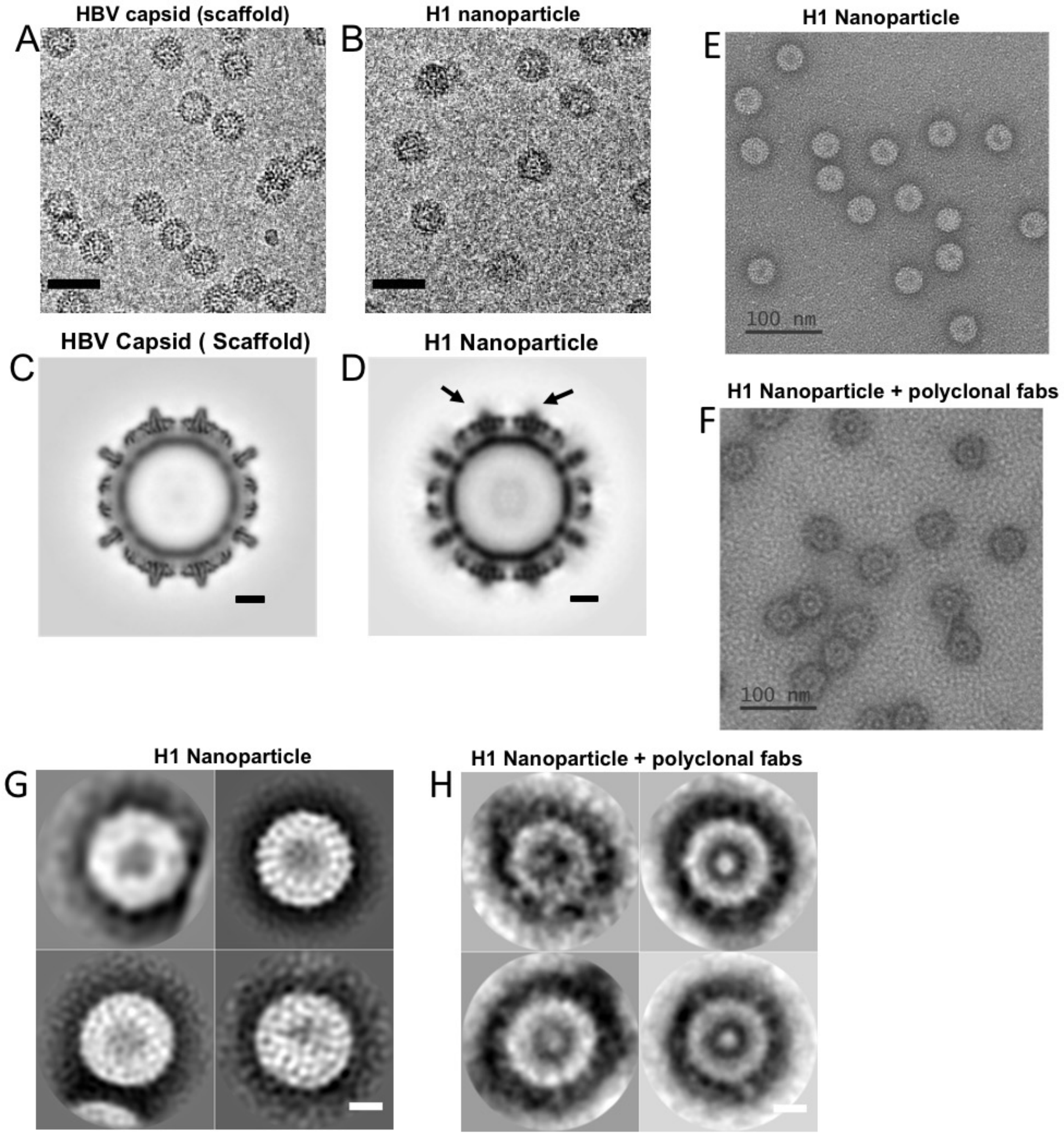

HBV capsid (scaffold)
A
H1 nanoparticle
B
H1 Nanoparticle
E
100 nm
HBV Capsid ( Scaffold)
C
H1 Nanoparticle
D
H1 Nanoparticle + polyclonal fabs
F
100 nm
H1 Nanoparticle
G
H1 Nanoparticle + polyclonal fabs
H


Fig. S8

A

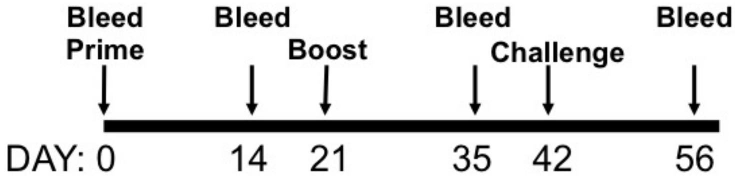


B

Fraction of Mouse Serum Samples able to elicit
Hemagglutination-Inhibition Antibody Titers (>10) to A/California/07/2009

| | Pre-Challenge | | | |
|---|---|---|---|---|
| | **Day 0** | **Day 14** | **Day 35** | **Day 56*** |
| **Saline** | 0/10 | 0/10 | 0/10 | 0/0 |
| **Nanoparticle** | 0/10 | 0/10 | 0/10 | 4/4 |
| **Nanoparticle + Adjuvant** | 0/10 | 0/10 | 0/10 | 2/2 |

*For day 56 samples only serum from challenge survivors were tested.

C

Fraction of Mouse Serum Samples able to elicit
Microneutralization Assay Titers (>20) to A/California/07/2009

| | Pre-Challenge | | | |
|---|---|---|---|---|
| | **Day 0** | **Day 14** | **Day 35** | **Day 56*** |
| **Saline** | 0/10 | 0/10 | 0/10 | 0/0 |
| **Nanoparticle** | 0/10 | 0/10 | 0/10 | 3/3 |
| **Nanoparticle + Adjuvant** | 0/10 | 0/10 | 0/10 | 2/2 |

*For day 56 samples only serum from challenge survivors were tested.

Fig. S9

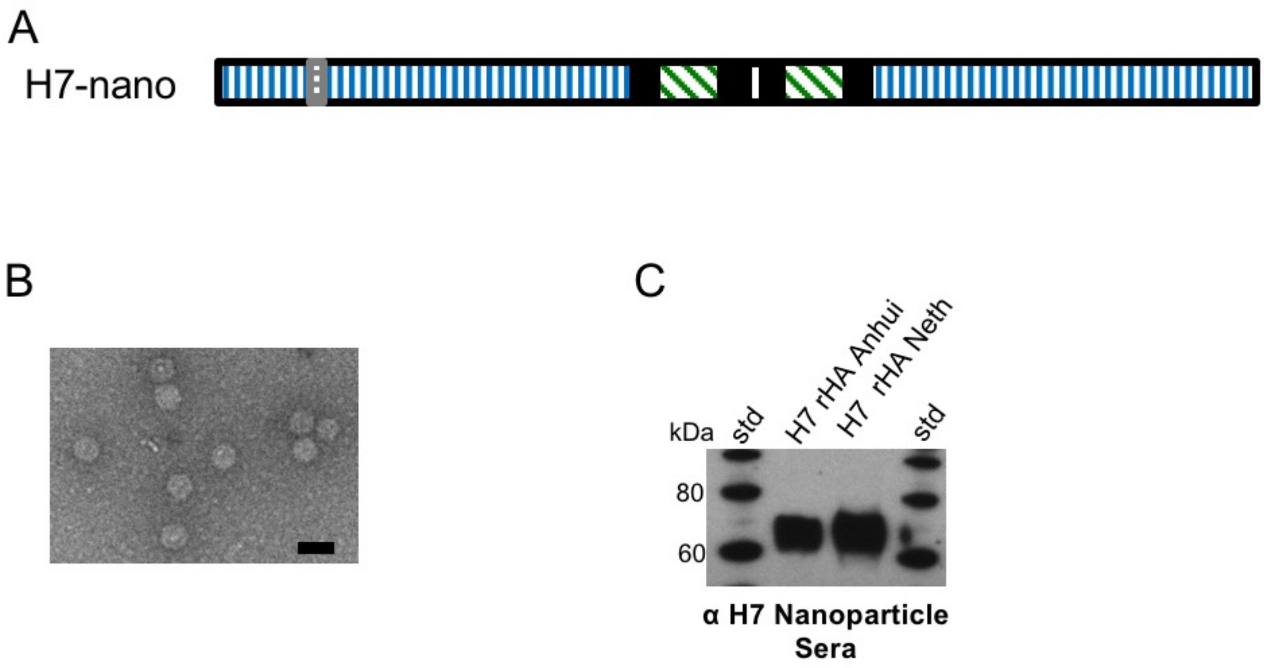
A
H7-nano
B
C
H7 rHA Anhui
H7 rHA Neth
kDa
std
std
80
60
α H7 Nanoparticle Sera

Fig. S10

**Supplemental Figure Captions.**

**Fig. S1. Footprints of broadly neutralizing stem antibodies.** (A-C) Structures of three broadly-reactive stem antibodies to influenza HA were colored to illustrate the extent of the bound Fab footprint on HA. HA1 is colored red, HA2 is colored blue, and the A-helix is colored green. Overlaid on the HA structure is the footprint for antibodies (A) C179, (B) CR6261, and (C) FI6v3, with respective PDB codes 4HLZ, 3GBN, and 3ZTJ. Footprints were depicted as HA atoms within 5 Å of Fab atoms, hydrogen atoms excluded. Footprints for each broadly neutralizing antibody extend beyond the A-helix residues. C179 is a mouse HA stem antibody while CR6261 and FI6v3 are human stem antibodies.

**Fig. S2. Bioinformatic analysis of helix A sequence conservation between hemagglutinin subtypes.** Hemagglutinin sequences (N=50,428) (H1 to H16) were downloaded from the influenza sequence database. Sequences were grouped according to subtype and the helix A region for each sequence was extracted. Consensus sequences for the helix A for each HA subtype was derived by comparison of all sequences within a group. Pair-wise sequence identity comparisons between different HA subtypes (H1-H16) were then organized as a matrix. Identities ranged from 50% to 100% for comparison between two different subtypes.

**Fig. S3. Strategy for the bioinformatic design of a nanoparticle library that displays helix-A for hemagglutinin (HA).** (A) Over 50,0000 sequences of HA from the influenza database were downloaded and curated into about 36,000 helix-A sequences for the 16 HA subtypes. A cDNA sequence library was created containing 27 HA helix A-nanoparticle sequences encompassing the HA subtypes. (B) Bioinformatic analysis of the helix A regions from the 27 HA-nanoparticle sequence library comparing sequence identity and coverage to the larger helix-A sequence database of over 36,000 HA sequences.

**Fig. S4. Schematics for construct designs and analysis of protein expression by immunoblotting.** (A) HBV capsid (scaffold) without helix-A insertions. The capsid sequence is represented by boxes with blue vertical lines. The immunodominant loop (c1 epitope) is represented as a white box and denoted by an asterisk. (B) Construct for H1-nanoparticle (H1-nano) consists of two copies of helix A of influenza hemagglutinin H1 HA (CA09) inserted into the scaffold at the immunodominant loop region to flank the c1 sequence on both sides . Each copy of helix-A is represented by a box with green diagonal lines. Each helix-A sequence is flanked by linker sequences (black boxes). The constructs contain an epitope tag for antibody 10E11 (gray box, horizontal lines) that recognizes residues 1-10 at the N-terminus of the capsid scaffold. (C) Diagram key to indicate the schematic representations for each region: HBV capsid scaffold, helix-A epitope insertion site into immunodominant lope (c1 epitope) of capsid scaffold, helix-A, linkers, and epitope tag. (D,E) Western blots to probe for expression of capsid scaffold (without helix-A) and H1-nanoparticle (with helix-A), respectively. Antibody 10E11 was used as primary antibody to detect the tag sequence. Molecular weight standards are denoted (std) and bands for scaffold and H1-nanoparticles are denoted with asterisks.

**Fig. S5. Ensemble of molecular models with variable positioning helix-A epitope on designed protein.** (A) Multiple molecular models of a dimeric unit of the H1-nanoparticle with the HBV capsid scaffold (gray) being fixed relative to the helix-A stem epitope having different conformations and orientations (green, red, gold yellow). In the scaffold monomers (gray) have two copies of HA helix A inserted into the tip of the loop of the capsid protein. (B). A molecular model for the H1-nanoparticle with helix-A stem epitopes in different positions as indicated in panel A. For clarity all helix-A portions are green with the scaffold in gray.

**Fig. S6. Purification and particle formation of scaffold and H1-nanoparticle constructs.** (A) SDS-PAGE analysis of purified HBV capsid (scaffold) purification fractions. The first lane contains molecular weight standards (std) and subsequent lanes contain gradient fractions for increasing sucrose concentrations. A white arrow indicates the bands at about 20 kDa. (B) Electron microscopy of purified HBV capsid (scaffold). Black arrows indicate capsid scaffold particles. Scale bar 50 nm. (C) SDS-PAGE analysis of H1-nanoparticle construct purification fractions. The first lane contains molecular weight standards (std) and subsequent lanes contain gradient fractions for increasing sucrose concentrations. A white arrow indicates the H1-nanoparticle protein bands at about 25 kDa. (D) Electron microscopy of purified H1-nanoparticles. Black arrows indicate H1-nanoparticles. Scale bar 50 nm.

**Fig. S7. Overview of H1 nanoparticle purification by centrifugation.** (A) Example of rotor and bucket used for gradient purification of nanoparticles. (B) Image of the H1-nanoparticle band in gradient, illuminated from below the tube. (C) Electron microscopy of H1-nanoparticles in the aforementioned gradient band after dialysis into PBS and (D) SDS-PAGE of the purified H1-nanoparticles.

**Fig. S8. Helix-A epitope disposition on H1-nanoparticle by structural comparisons of HBV capsid (scaffold) and H1 nanoparticles by electron microscopy.** (A) Cryo-electron microscopy images of HBV capsid (scaffold) particles and (B) H1-nanoparticles. (C) 3D reconstruction cross-section for HBV capsid scaffold by cryo-electron microscopy. Scale bar is 5 nm. (D) 3D reconstruction cross-section for H1-nanoparticles by cryo-electron microscopy. Arrows denote the location of the insertion, where diffuse density is observed compared to the HBV capsid scaffold. Scale bar is 5 nm. H1-nanoparticle subunits that comprise the nanoparticle give the appearance of spikes on the surface. (E) Negative-stain electron microscopy of H1-nanoparticles incubated without Fabs and (F) incubated with Fabs derived from polyclonal sera from mice immunized with H1-nanoparticles. Scale bar is 100 nm. (G) Images of 2D class-averages from 591 PTA stained H1-nanoparticles classified into 4 classes. The best resolved class (upper-right) illustrates a protein shell, measures about 7 nm thick, and encapsulates a slightly darker core. H1-nanoparticle subunits that comprise the nanoparticle give the appearance of undulations of the particle surface when visualized by PTA stain. (H) Images of 2D class averages from 308 PTA stained H1-nanoparticles in complex with polyclonal Fabs were classified into 4 classes. The extra density surrounding the H1-nanoparticles, visible in all 4 averages, is consistent with a coat of bound polyclonal Fab. Scale bars are 10 nm. Note, for ease of comparison between HBV capsid (scaffold) structure and H1-nanoparticle, panel D is the same as main Figure 2E.

**Fig. S9. Hemagglutinin-inhibition and neutralization activity for mouse sera after immunization with HA nanoparticles.** (A) Schedule for mouse immunization and challenge. Mice were immunized on days 0 and 21 with H1-nanoparticle with and without adjuvant. Mice were challenged on day 42 with 10X Mouse Lethal Dose ($MLD_{50}$) of H1N1 virus. (B) Time course of hemagglutination-inhibition activity (HAI) for mouse sera. (C) Time course of microneutralization (MN) activity for mouse sera. Greater than 10 HAI and >20 MN were chosen because these were above judged background levels of the assays. Since the sera were taken pre-challenge the denominators are 10 and for day 56. Only serum from challenge survivors were tested and thus there are lower denominators. Note, for ease of comparison S9A is the same immunization figure as main figure 5A. Saline is PBS. Nanoparticles is H1-nanoparitcle and adjuvant is SAS (Sigma Adjuvant System, oil-in-water emulsion).

**Fig. S10. Design, purification, and immunogenicity of H7 nanoparticle.** (A) Schematic for construct design for H7-nanoparticle. The helix-A HA-nanoparticle (H7-nano) construct consists of two copies of helix A of influenza H7 HA inserted into the HBV capsid scaffold at the immunodominant loop region. Each copy of helix-A is represented by a box with green diagonal lines. Each helix-A sequence is flanked by linker sequences (black boxes). The construct contains an epitope tag for antibody 10E11 (gray box, horizontal lines) that recognizes residues 1-10 at the N-terminus of the capsid scaffold. (B) Purified H7-nanoparticles observed by negative-stained electron microscopy. Scale bar, 50 nm. (C) Western blot displaying reactivity of H7-nanoparticle mice sera to recombinant H7 HA proteins from H7N9 (H7 rHA Anhui), and H7N7 (H7 rHA Netherland [Neth]) viruses.
